# Cytogenotoxic effects of 3-epicaryoptin in *Allium cepa* L. root apical meristem cells

**DOI:** 10.1101/2021.03.26.437299

**Authors:** Manabendu Barman, Sanjib Ray

**Affiliations:** Molecular Biology and Genetics Unit, Department of Zoology, The University of Burdwan, Golapbag, Purba Bardhaman-713104, West Bengal, India

**Keywords:** 3-epicaryoptin, *Allium cepa* test, chromosomal abnormalities, micronuclei, polyploidy

## Abstract

Diterpenoid 3-epicaryoptin (C_26_H_36_O_9_) is abundant in the leaves of *Clerodendrum inerme*, a traditionally used medicinal plant, having insect antifeedant activities. Here, we aim to explore the cytogenotoxic effects of compound 3-epicaryoptin in *Allium cepa* root apical meristem cells. *A. cepa* roots were treated with 3-epicaryoptin (100, 150 & 200 μg mL^-1^ concentration) and the standard compound colchicine (200 μg mL^-1^ concentration) for 2, 4, 4+16 h (4 h treatment followed by 16 h recovery). Cytogenotoxicity was analysed by studying the root growth retardation (RGR), mitotic index (MI), and cellular aberrations. The result showed statistically significant (*p*<0.01), concentration-dependent RGR effects of 3-epicaryoptin treatment compared with the negative control. Study of cell frequency in different phases of cell division observed a significant (*p*<0.001) increase in the metaphase cell percentage (66.2±0.58 %, 150 μg mL^-1^) and which subsequently caused an increase in the frequency of MI (12.29±0.34 %, 150 μg mL^-1^) at 4h of 3-epicaryoptin treatment and that was comparable with the colchicine action. The cytological study revealed that the 3-epicaryoptin treatment could induce different types of chromosomal abnormalities such as colchicine like metaphase, vagrant chromosomes, sticky chromosomes, anaphase-bridge, and an increased frequency of micronuclei and polyploid cells. These findings indicate that 3-epicaryoptin is cytogenotoxic, and thus *C. inerme* should be used with caution in traditional medicine.

## Introduction

3-epicaryoptin is a neo-clerodane diterpenoid compound found in a few species of the *Clerodendrum* genus, such as *C. calamitosum, C. paniculatum*, and *C. inerme*. Besides these, it has also been reported to be present in the genus *Ajuga* (*A. bracteosa*) (Hosozawa et al. 1974; Pereira and Gurudutt 1990; Krishna kumari et al. 2003; Castro et al. 2011). It was first isolated and identified by Hosozawa et al. (1974) from the leaves of *C. calamitosum* Maxim (Verbenaceae) (Hosozawa et al. 1974). The previous study reports showed that 3-epicaryoptin possesses potent insect antifeedant activity against the third instar larvae of the tobacco cutworm, *Spodoptera litura* F. and fourth instar nymphs and adults of the potato beetle *Henosepilachna vigntioctopunctata* Fab. (Coleoptera: Coccinellidae) (Hosozawa et al. 1974; Govindachari et al. 1998). In addition to insect antifeedant activity, 3-epicaryoptin reduced the growth and increased the mortality of *Ostrinia nubilalis* (Hubner) (European com borer) larvae (Beninger et al. 1993). It also inhibited the development of *Musca domestica* and *Culex quinquefasciatus* larvae (Pereira and Gurudutt 1990). Although the insect antifeedant and larvicidal potential of 3-epicaryoptin has been well documented, yet there is no report available on its cytogenotoxic effects.

*Allium cepa* L. is one of the most established plant bioassays for cytotoxicological assessment due to its kinetic proliferation properties, large chromosomes, low chromosome number (2*n* = 16), easily observed with a light microscope, as validated by the UNEP and the IPCS (Cabrera and Rodriguez 1999; Gomes et al. 2013). The *A. cepa* test system provides important information about the possible mechanisms of action of agents on the genetic material and has a strong correlation with mammalian/cell culture systems (Leme and Marin-Morales 2009; Fiskesjö 1985; Rank and Nielsen 1994; Chauhan et al. 1999; Tedesco and Laughinghouse 2012). The use of *A. cepa* as a test system was first investigated by Levan, demonstrating disturbances in the mitotic spindle apparatus due to the use of colchicine (Levan 1938). Studies have found that colchicine treatment causes root growth inhibition and swelling effects, increase the frequency of metaphase cells with haphazardly arranged condensed chromosomes by inhibiting mitotic spindle organization. As a result of this, chromatids fail to move to the opposite poles and eventually become enclosed in a new nuclear membrane and proceed into interphase as a doubled chromosome number of polyploid (PP) cells (Hague and Jones 1987). Treatment of colchicine in *A. cepa* could also induce an increased frequency of chromosomal abnormalities (CA) and micronucleated (MN) cells (Ray et al. 2013). In this study, colchicine was used as a positive control to determine the cytogenotoxicity of compound 3-epicaryoptin in *A. cepa* root tip cells.

In the previous study, we found that the leaf aqueous extract of *C. inerme* (LAECI) has colchicine like *A. cepa* root tip swelling (RTS) effects and increases the frequencies of colchicine-like metaphase (c-metaphase) cells together with the formation of other chromosomal abnormalities, MN, and polyploidy (PP) (Barman et al. 2020; 2021; Barman and Ray 2022). However, the active compound responsible for this activity has not been well studied. Given these findings, the focus of the current work was to evaluate the cytogenotoxicity of compound 3-epicaryoptin using the *A. cepa* assay.

## Materials and methods

### Chemicals

Glacial acetic acid, orcein, and methanol were purchased from Merck Ltd. Mumbai, India. Colchicine was obtained from Himedia Laboratories Pvt. Ltd. Mumbai, India. Compound 3-epicaryoptin was isolated from *C. inerme* leaf aqueous extract (Ray et al. 2019).

### Effects of 3-epicaryoptin on *Allium cepa*

*Allium cepa* L. root apical meristem cells were used as a plant model for determining the cytogenotoxicity of compound 3-epicaryoptin by the study of root growth inhibitory effects, MI, CA, MN, and PP.

### Root growth retardation and swelling effects

The similar sized *A. cepa* bulbs were purchased from a local vegetable store at Golapbag, Burdwan, and allowed for root sprouting as described earlier (Barman et al. 2020; Ray et al. 2013). For root growth inhibition analysis, the sprouting roots were exposed continuously to the different concentrations of 3-epicaryoptin (12.5, 25, 50, 100, 200 μg mL^-1^) and colchicine (200 μg mL^-1^) for 24, 48 and 72 h. The root growth inhibition percentage and the IC_50_ value were determined at 24, 48 and 72 h. 3-epicaryoptin (100, 150 and 200 μg mL^-1^) and standard colchicine (200 μg mL^-1^) induced morphometric RTS effects were also measured after 4 h treatment followed by16 h recovery in water (4+16 h).

### Mitotic index and cellular aberrations studies

The onion roots having 1.5-2 cm in length were exposed to the different concentrations (100, 150, and 200 μg mL^-1^) of 3-epicaryoptin for 2, 4, and 4+16 h. The effects were compared with the standard spindle poison, colchicine (200 μg mL^-1^). In the case of control groups, the onion roots were maintained in distilled water. For root tip squash preparation, the control and treated onion root tips were fixed in aceto-methanol 3(methanol): 1(acetic acid) for 24 h, then hydrolysed in 1N HCl at 60°C for 10 min, stained with aceto-orcein (2%) and finally squashed in 45% acetic acid (Chaudhuri and Ray 2015). The squashed root tips were observed in a bright field light microscope at 400X magnification. Digital images were taken with Future Win Joe, Future Optics (Version 1.6.5.1207), and then calculating the frequency of MI, dividing cell percentage at different phases (prophase, metaphase, anaphase, telophase), as well as cellular abnormalities, i.e. CA, MN, and PP cells, from the micrographs.

### Scoring and statistical analysis

Data obtained on MI, (MI (%) = Number of dividing cells/Total number of cells scored X 100), cell percentage at different phases, CA, MN, and PP cell frequencies were analysed by using the 2 × 2 contingency χ^2^ test, and the IC_50_ value for the RGR was calculated with probit analysis in SPSS Version 20. Differences between control and treated groups were considered as significant at *p*≤ 0.05 or 0.01 or 0.001. All the data were expressed as mean ± SEM (standard error of mean). Correlations among dividing phase and among different cellular abnormalities were calculated using Pearson’s bivariate correlations and analyses were done using PAST version 4.05.

## Results

### Root growth retardation and swelling effects

Treatment of 3-epicaryoptin causes concentration-dependent, statistically significant RGR effects in *A. cepa* root apical meristem cells (**Fig. 1**). The maximum RGR (89.77%, *p*< 0.001) effect of 3-epicaryoptin treatment was at 200 μg mL^-1^ concentration at 72 h. The standard colchicine (200 μg mL^-1^) induced the highest RGR percentage, 92.08% (*p*< 0.001) at 72 h. The IC_50_ values were 68.38, 70.50, and 63.96 μg mL^-1^ respectively, at 24, 48, and 72 h of 3-epicaryoptin treatment. The RTS phenomenon was also observed in 3-epicaryoptin (100, 150, and 200 μg mL^-1^) and colchicine (200 μg mL^-1^) treated samples after 4+16 h. A comparatively better RTS effect was observed at a concentration of 200 μg mL^-1^ of 3-epicaryoptin than with colchicine treatments (200 μg mL^-1^) (**Fig. 1**).

**Fig. 1.**
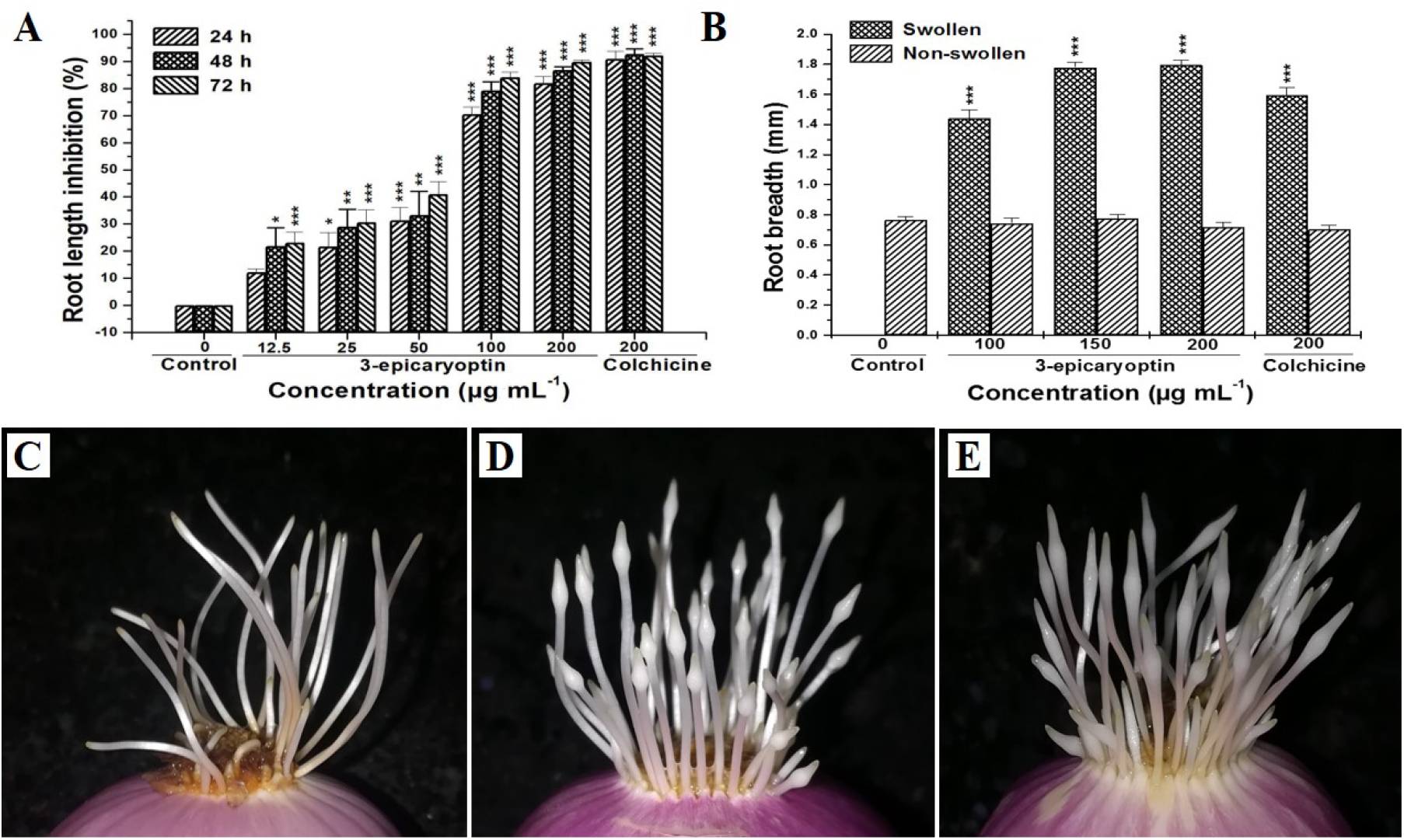
The effects of 3-epicaryoptin and colchicine in the *A. cepa* root length inhibition % after 24, 48, and 72 h continuous treatment (A), and the root tip swelling effects at 4+16 h treatment (B-E). (B) Measurement the diameter of the *A. cepa* root tips, (C) Control, (D) 200 μg mL^-1^ concentration of 3-epicaryoptin and (E) 200 μg mL^-1^ concentration of colchicine induced *A. cepa* root tip swelling effects.

### Effect on mitotic index and dividing phase frequency

The effects of 3-epicaryoptin and colchicine on the MI% and dividing phases in *A. cepa* root apical meristem cells are presented in **Table 1**. Data indicate that treatment with 3-epicaryoptin (100, 150, and 200 μg mL^-1^) and colchicine (200 μg mL^-1^) causes an increase in MI% (*p*< 0.001) at 2 and 4 h treated onion root tip cells. However, in the case of 16 h water recovery treatment, a dose-dependent decrease in MI% was observed as compared to the untreated controls. Here, 3-epicaryoptin induced the highest MI% at 150 μg mL^-1^ (12.29%) at 4 h, whereas at 16 h recovery, it was decreased to 4.90% (*p*< 0.01). In the case of 200 μg mL^-1^ colchicine treatment, a statistically significant increase in MI % (13.18%, *p*< 0.001) was also observed at 4 h and again decreased (3.22%, *p*< 0.01) at 16 h recovery samples. Study of dividing phase frequency revealed that 3-epicaryoptin (150 and 200μg mL^-1^) continuous treatment for 4 h causes a significant decrease in prophase % (23.22% and 24.49%) and increase in metaphase % (66.2% and 63.31%) when compared to control (28.84%). The standard colchicine (200μg mL^-1^) at 4 h also decreased prophase (11.3%) and increased (82.35%) the metaphase cell percentage. Both 3-epicaryoptin and colchicine treatments increased the prophase-metaphase cumulative frequency (PMCF) and significantly decreased the anaphase-telophase cumulative frequency (ATCF) (**Fig. 2**). However, both PMCF and ATCF were always significantly higher in the case of colchicine-treated samples. The increasing tendency of PMCF was observed at 4 h and scored as 71.43±0.59%, 83.03±0.59%, 89.43±0.87%, and 87.81±0.26% respectively, for 0, 100, 150, and 200 μg mL^-1^ of 3-epicaryoptin and 93.66±1.35 % for 200 μg mL^-1^ of colchicine treatments. On the other hand, at 4 h treatment, decreasing tendency of ATCFs were scored as 28.78±0.78, 16.95±0.59, 10.55±0.87, and 12.17±0.26% respectively for 0, 100, 150, and 200 μg mL^-1^ of 3-epicaryoptin and 6.28±0.73% for 200 μg mL^-1^of colchicine.

**Table 1.**
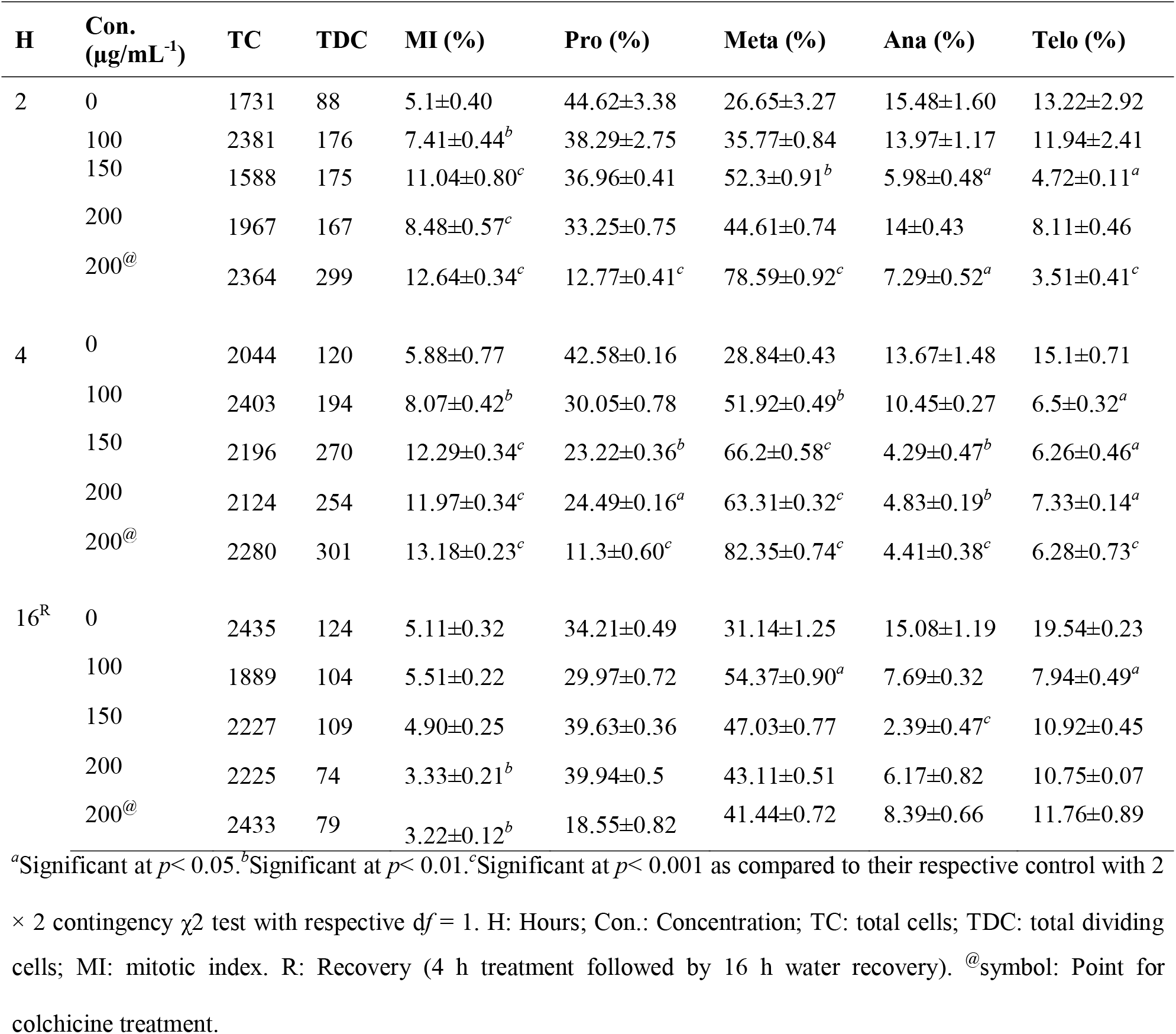
Influence of 3-epicaryoptin and colchicine on MI and percentages of the different cell division phases of *A. cepa* root apical meristem cells.

| H | Con.<br>($\mu\text{g/mL}^{-1}$) | TC | TDC | MI (%) | Pro (%) | Meta (%) | Ana (%) | Telo (%) |
| --- | --- | --- | --- | --- | --- | --- | --- | --- |
| 2 | 0 | 1731 | 88 | $5.1 \pm 0.40$ | $44.62 \pm 3.38$ | $26.65 \pm 3.27$ | $15.48 \pm 1.60$ | $13.22 \pm 2.92$ |
| | 100 | 2381 | 176 | $7.41 \pm 0.44^b$ | $38.29 \pm 2.75$ | $35.77 \pm 0.84$ | $13.97 \pm 1.17$ | $11.94 \pm 2.41$ |
| | 150 | 1588 | 175 | $11.04 \pm 0.80^c$ | $36.96 \pm 0.41$ | $52.3 \pm 0.91^b$ | $5.98 \pm 0.48^a$ | $4.72 \pm 0.11^a$ |
| | 200 | 1967 | 167 | $8.48 \pm 0.57^c$ | $33.25 \pm 0.75$ | $44.61 \pm 0.74$ | $14 \pm 0.43$ | $8.11 \pm 0.46$ |
| | 200<sup>@</sup> | 2364 | 299 | $12.64 \pm 0.34^c$ | $12.77 \pm 0.41^c$ | $78.59 \pm 0.92^c$ | $7.29 \pm 0.52^a$ | $3.51 \pm 0.41^c$ |
| 4 | 0 | 2044 | 120 | $5.88 \pm 0.77$ | $42.58 \pm 0.16$ | $28.84 \pm 0.43$ | $13.67 \pm 1.48$ | $15.1 \pm 0.71$ |
| | 100 | 2403 | 194 | $8.07 \pm 0.42^b$ | $30.05 \pm 0.78$ | $51.92 \pm 0.49^b$ | $10.45 \pm 0.27$ | $6.5 \pm 0.32^a$ |
| | 150 | 2196 | 270 | $12.29 \pm 0.34^c$ | $23.22 \pm 0.36^b$ | $66.2 \pm 0.58^c$ | $4.29 \pm 0.47^b$ | $6.26 \pm 0.46^a$ |
| | 200 | 2124 | 254 | $11.97 \pm 0.34^c$ | $24.49 \pm 0.16^a$ | $63.31 \pm 0.32^c$ | $4.83 \pm 0.19^b$ | $7.33 \pm 0.14^a$ |
| | 200<sup>@</sup> | 2280 | 301 | $13.18 \pm 0.23^c$ | $11.3 \pm 0.60^c$ | $82.35 \pm 0.74^c$ | $4.41 \pm 0.38^c$ | $6.28 \pm 0.73^c$ |
| 16<sup>R</sup> | 0 | 2435 | 124 | $5.11 \pm 0.32$ | $34.21 \pm 0.49$ | $31.14 \pm 1.25$ | $15.08 \pm 1.19$ | $19.54 \pm 0.23$ |
| | 100 | 1889 | 104 | $5.51 \pm 0.22$ | $29.97 \pm 0.72$ | $54.37 \pm 0.90^a$ | $7.69 \pm 0.32$ | $7.94 \pm 0.49^a$ |
| | 150 | 2227 | 109 | $4.90 \pm 0.25$ | $39.63 \pm 0.36$ | $47.03 \pm 0.77$ | $2.39 \pm 0.47^c$ | $10.92 \pm 0.45$ |
| | 200 | 2225 | 74 | $3.33 \pm 0.21^b$ | $39.94 \pm 0.5$ | $43.11 \pm 0.51$ | $6.17 \pm 0.82$ | $10.75 \pm 0.07$ |
| | 200<sup>@</sup> | 2433 | 79 | $3.22 \pm 0.12^b$ | $18.55 \pm 0.82$ | $41.44 \pm 0.72$ | $8.39 \pm 0.66$ | $11.76 \pm 0.89$ |

<sup>a</sup>Significant at $p < 0.05$ . <sup>b</sup>Significant at $p < 0.01$ . <sup>c</sup>Significant at $p < 0.001$ as compared to their respective control with 2 × 2 contingency $\chi^2$ test with respective $df = 1$ . H: Hours; Con.: Concentration; TC: total cells; TDC: total dividing
cells; MI: mitotic index. R: Recovery (4 h treatment followed by 16 h water recovery). @symbol: Point for colchicine treatment.

**Fig. 2.**
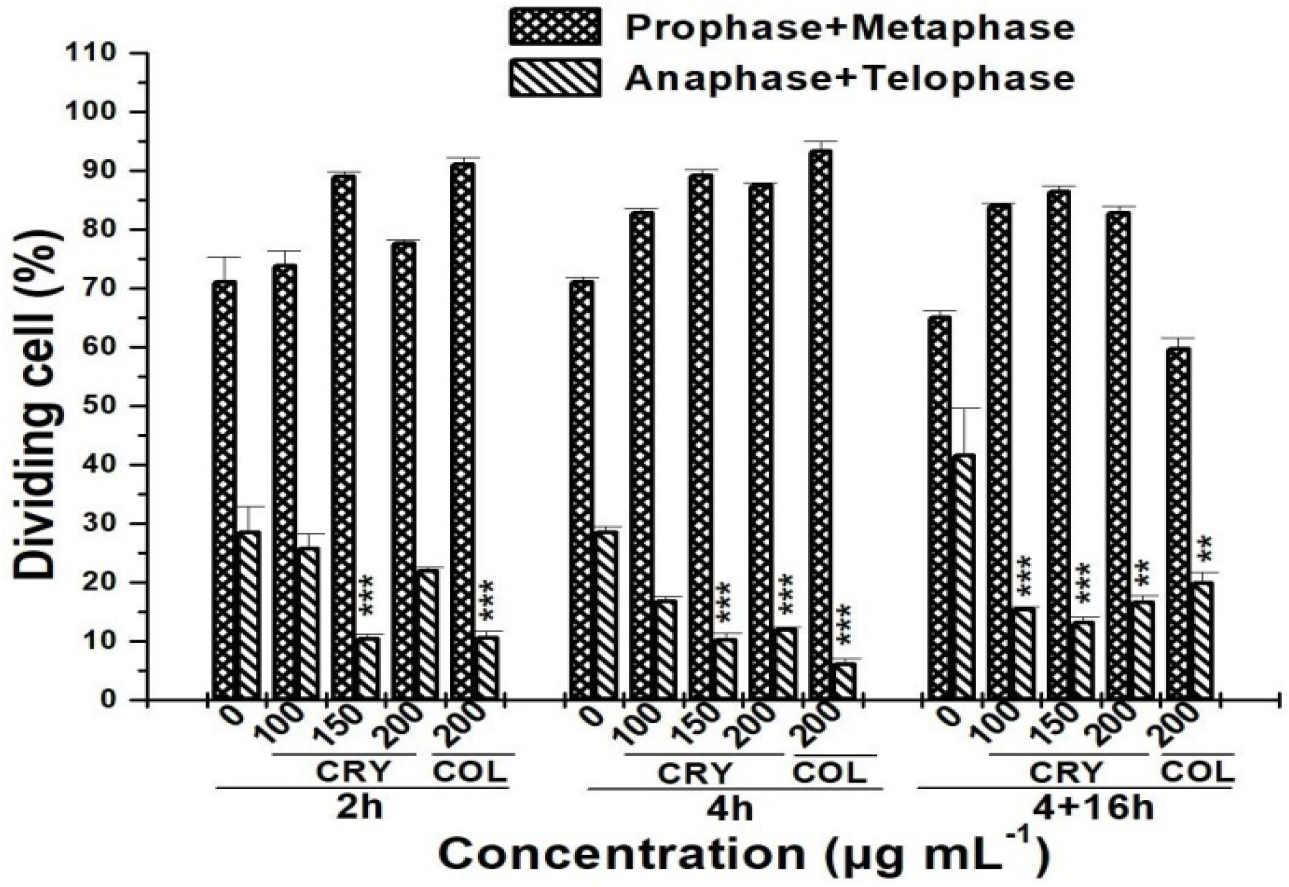
Influence of 3-epicaryoptin (100, 150, 200 μg mL^-1^) and colchicine (200 μg mL^-1^) on the ‘prophase-metaphase’ and ‘anaphase-telophase’ cumulative frequencies in *A. cepa* root tip cells. *Significant at *p*< 0.05, \*\**p*< 0.01 and \*\*\**p* < 0.001 with 2 × 2 contingency χ^2^ analysis compared to respective control at *df* = 1. CRY: 3-epicaryoptin; COL: Colchicine.

### Chromosomal abnormalities

3-epicaryoptin induced the formation of different types of CA in *A. cepa* root meristem cells and which are comparable to colchicine effects (**Table 2 and Fig. 3**). 150 μg mL^-1^ of 3-epicaryoptin treatment showed a significant (*p*< 0.001) increase in the total CA% (8.79%, 7.8%, and 30.46%) as compared to standard colchicine (13.48%, 15.91%, and 33.48%) respectively at 2, 4, and 16 h water recovery treatment. The most visible CA induced by 3-epicaryoptin was c-metaphase. The mean percentages of c-metaphase were 29.61±0.81%, 51.56±0.57%, and 41.04±0.64% at 2 h (*p*< 0.001) and 40.95±0.87%, 63.39±0.51%, and 60.43±0.57 % at 4 h (*p*< 0.001) for the respective concentration of 100, 150, and 200 μg mL^-1^ of 3-epicaryoptin, whereas, the negative control showed 1.01±0.50% and 0.45±0.45% of c-metaphase cells at the respective fixative hours. While 200 μg mL^-1^ colchicine showed the highest c-metaphase frequency (72.65±0.11 % and 77.24±0.10 %) at both 2 h (*p* < 0.001) and 4 h (*p*< 0.001) treated samples (**Table 2**). Treatment with 3-epicaryoptin also increased the frequency of vagrant chromosomes. 150 μg mL^-1^ of 3-epicaryoptin treatment was found to be 13.89±0.21% (*p*<0.01), 13.64±0.34% (*p*<0.01), and 15.09±0.49 % (*p*<0.001) of vagrant chromosome after 2, 4, and 16 h recovery in water. Colchicine (200 μg mL^-1^) induced the highest vagrant chromosome percentage, which was observed to be 11.53±0.03% at 4 h of treatment. A significant frequency (*p*< 0.05) of sticky chromosomes was also observed at 150 μg mL^-1^ (12.13±0.54 %) and 200 μg mL^-1^ (11.27±0.19 %) of 3-epicaryoptin and 200 μg mL^-1^ (15.15±0.29 %) of colchicine treatment after 4 h. Besides the above abnormalities, 3-epicaryoptin and colchicine could increase the frequency of anaphase-bridge, lagging chromosome, polar deviation, and multipolar anaphase-telophase. The frequency was not statistically significant as compared to the respective control group (**Table 2 and Fig. 3**).

**Table 2.** Effects of 3-epicaryoptin and colchicine on the frequency of CA in *A. cepa* root apical meristem cells.

| H | Conc.<br>($\mu\text{g mL}^{-1}$) | TC | TDC | % of AB<br>cell | Ana-Bridge<br>(%) | C-meta (%) | Vagrant (%) | Cr Sti (%) | Lagging<br>Cr (%) | Polar<br>deviation<br>(%) | Multipolar<br>Ana Telo<br>(%) |
| --- | --- | --- | --- | --- | --- | --- | --- | --- | --- | --- | --- |
| 2 | 0 | 1245 | 66 | $0.34\pm0.01$ | $1.01\pm0.5$ | $1.01\pm0.5$ | $0.54\pm0.54$ | $0.46\pm0.46$ | $0.95\pm0.47$ | 0 | 0 |
| | 100 | 1160 | 78 | $3.53\pm0.36^c$ | $7.66\pm0.15$ | $29.61\pm0.81^c$ | $10.63\pm0.29$ | $8.04\pm0.48$ | $8.59\pm0.37$ | $3.08\pm0.67$ | $3.4\pm0.27$ |
| | 150 | 1175 | 132 | $8.79\pm0.01^c$ | $8.28\pm0.62$ | $51.56\pm0.57^c$ | $13.89\pm0.21^b$ | $10.25\pm0.33$ | $4.23\pm0.35$ | $1.37\pm0.24$ | $3.25\pm0.36$ |
| | 200 | 1356 | 114 | $5.25\pm0.40^c$ | $7.08\pm0.34$ | $41.04\pm0.64^c$ | $9.85\pm0.56$ | $9.16\pm0.48$ | $4.91\pm0.25$ | $0.78\pm0.43$ | $4.91\pm0.25$ |
| | 200<sup>@</sup> | 1453 | 185 | $13.48\pm0.26^c$ | $4.85\pm0.02$ | $72.65\pm0.11^c$ | $9.70\pm0.04$ | $11.68\pm0.30^a$ | $1.25\pm0.17$ | $0.53\pm0.00$ | $1.61\pm0.00$ |
| 4 | 0 | 1287 | 71 | $0.25\pm0.02$ | $0.88\pm0.44$ | $0.45\pm0.45$ | $0.97\pm0.48$ | $0.52\pm0.52$ | $0.88\pm0.44$ | $0.52\pm0.52$ | 0 |
| | 100 | 1306 | 88 | $4.69\pm0.09^c$ | $7.24\pm0.53$ | $40.95\pm0.87^c$ | $9.07\pm0.09$ | $10.26\pm0.56$ | $11.28\pm0.6$ | $2.34\pm0.79$ | $5.35\pm0.56$ |
| | 150 | 1252 | 127 | $7.8\pm0.08^c$ | $9.91\pm0.34$ | $63.39\pm0.51^c$ | $13.64\pm0.34^b$ | $12.13\pm0.54^a$ | $6.79\pm0.05$ | $1.53\pm0.36$ | $11.26\pm0.24$ |
| | 200 | 1419 | 148 | $8.97\pm0.61^c$ | $4.91\pm0.27$ | $60.43\pm0.57^c$ | $12.36\pm0.18^b$ | $11.27\pm0.19^a$ | $4.68\pm0.48$ | $0.88\pm0.16$ | $6.96\pm0.15$ |
| | 200<sup>@</sup> | 1520 | 211 | $15.91\pm0.54^c$ | $3.68\pm0.26$ | $77.24\pm0.10^c$ | $11.53\pm0.03^b$ | $15.15\pm0.29^b$ | $0.47\pm0.00$ | 0 | $1.92\pm0.00$ |
| 16<sup>R</sup> | 0 | 1176 | 63 | $0.27\pm0.01$ | $0.49\pm0.49$ | $1.59\pm0.09$ | $1\pm0.5$ | $0.59\pm0.59$ | $0.50\pm0.50$ | 0 | 0 |
| | 100 | 1161 | 64 | $19.95\pm0.78^c$ | $3.13\pm0.14$ | $23.36\pm0.75^c$ | $6.75\pm0.28$ | $6.27\pm0.28$ | $4.7\pm0.21$ | $1.56\pm0.07$ | $6.75\pm0.28$ |
| | 150 | 1085 | 26 | $30.46\pm0.63^c$ | $7.54\pm0.24$ | $35.57\pm0.29$ | $15.09\pm0.49$ | $8.38\pm0.84$ | $0.83\pm0.83$ | $0.83\pm0.83$ | $7.54\pm0.24$ |
| | 200 | 1229 | 34 | $26.41\pm0.61^c$ | $3.9\pm0.66$ | $21.37\pm0.73$ | $11.77\pm0.50$ | $5.63\pm1.01$ | $1.73\pm0.86$ | $0.83\pm0.83$ | $4.8\pm0.41$ |
| | 200<sup>@</sup> | 1505 | 50 | $33.48\pm0.44^c$ | $1.22\pm0.62$ | $5.81\pm0.53$ | $1.47\pm0.74$ | $2.56\pm0.36$ | 0 | $0.54\pm0.05$ | $5.26\pm0.43$ |

<sup>a</sup>Significant at $p<0.05$ , <sup>b</sup>Significant at $p<0.01$ , <sup>c</sup>Significant at $p<0.001$ as compared to their respective control with
$2 \times 2$ contingency $\chi^2$ test with respective $df = 1$ . TC: total cells; H: Hours; TC: Total Cells; TDC: total dividing cells; AB: Aberrant; Meta: Metaphase; Ana: Anaphase; Telo: Telophase; Cr: Chromosome; Sti: Stickiness. R: Recovery (4 h treatment followed by 16 h water recovery); <sup>@</sup>symbol: Point for colchicine treatment.

**Fig. 3.**
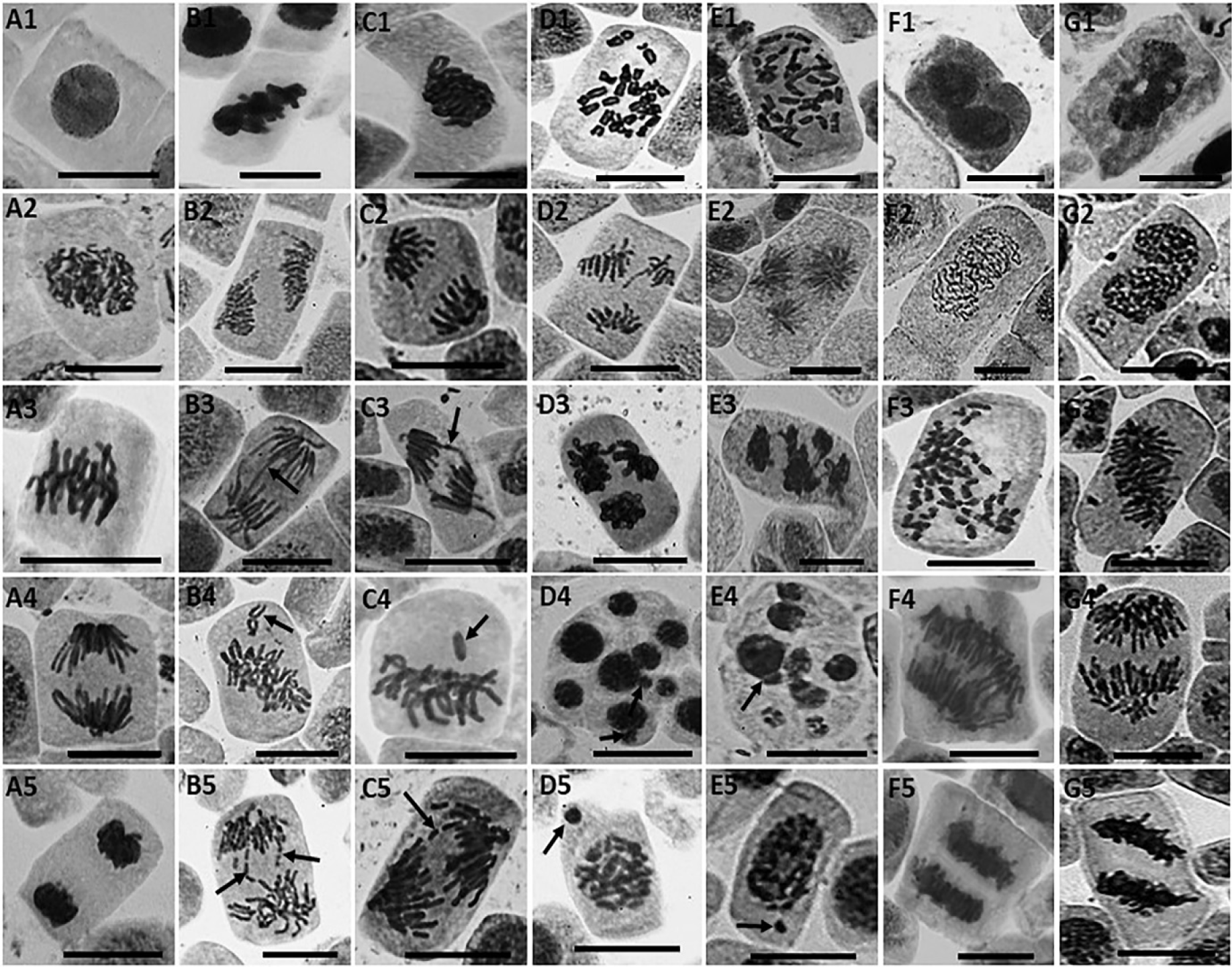
Shows 3-epicaryoptin and colchicine induced formation of different types of CA, MN and PP cells in *A. cepa* root apical meristem cells. (A1-A5) control interphase, prophase, metaphase, anaphase and telophase; (B1-B5) 3-epicaryoptin induced formation of: (B1) Sticky chromosome at metaphase (150 μg mL^-1^/4 h), (B2) Polar deviation at anaphase (100 μg mL^-1^/16 h R), (B3) Anaphase bridge (200 μg mL^-1^/2 h), (B4) Vagrant chromosome at metaphase (100 μg mL^-1^/4 h), (B5) Lagging chromosome at anaphase (200 μg mL^-1^/2 h); (C1-C5) 200 μg mL^-1^ colchicine induced formation of: (C1) Sticky chromosome at metaphase (4 h), (C2) polar deviation at anaphase (2 h), (C3) Anaphase bridge (2 h), (C4) Vagrant chromosome at metaphase (16 h R), (C5) Lagging chromosome at anaphase (4 h); (D1-D2) 3-epicaryoptin induced formation of: (D1) C-metaphase (100 μg mL^-1^/4 h), (D2) Multipolar anaphase (200 μg mL^-1^/4 h), (D3) Multipolar telophase (150 μg mL^-1^/2 h) (D4) cells with multiple MN and nuclear buds (150 μg mL^-1^/16 h R), (D5) MN at prophase (100 μg mL^-1^/16 h R); (E1-E5) 200 μg mL^-1^ colchicine induced formation of: (E1) C-metaphase (4 h), (E2) Multipolar anaphase (16 h R), (E3) Multipolar telophase (16 h R), (E4) cells with multiple MN and nuclear buds (16 h R), (E5) MN at prophase (4+16 h); (F1-F5) 3-epicaryoptin induced formation of: (F1) PP interphase (150 μg mL^-1^/4+16 h) (F2) PP prophase (100 μg mL^-1^/16 h R (F3) PP metaphase (100 μg mL^-1^/16 h R), (F4) PP anaphase (200 μg mL^-1^/16 h R) and (F5) PP telophase (150 μg mL^-1^); (G1-G5) 200 μg mL^-1^ colchicine induced formation of: (G1) PP interphase (16 h R) (G2) PP prophase (16 h R), (G3) PP metaphase (16 h R), (G4) PP anaphase (16 h R) and (G5) PP telophase (16 h R). R; Recovery (4 h treatment followed by recovery).

### Micronuclei and polyploidy

Both 3-epicaryoptin and colchicine treatments induced the formation of MN and PP cell in *A. cepa* root apical meristem (**Fig. 4 and 5**). A significant increase in the frequency of MN and PP cells was observed at 4 h treatment and also at 4+16 h water recovery. The MN frequencies were 12.94±0.34, 24.84±0.54, and 22.10±0.73% respectively, for 100, 150, and 200 μg mL^-1^ of 3-epicaryoptin treated samples after 4+16 h (*p*< 0.001). While, 200 μg mL^-1^ colchicine showed the highest MN percentage, 27.73±0.6%, at 16 h (*p*<0.001) recovery treatment (**Fig. 4**). 3-epicaryoptin treatment also statistically increased (*p*<0.001) the PP cell frequency at 4+16 h. The 150 μg mL^-1^ concentration of 3-epicaryoptin showed 30.61±0.6% increase in the PP cell frequency. Whereas the standard colchicine (200 μg mL^-1^) induced 32.66±0.79% increase in PP cell frequency (**Fig. 5**).

**Fig. 4.**
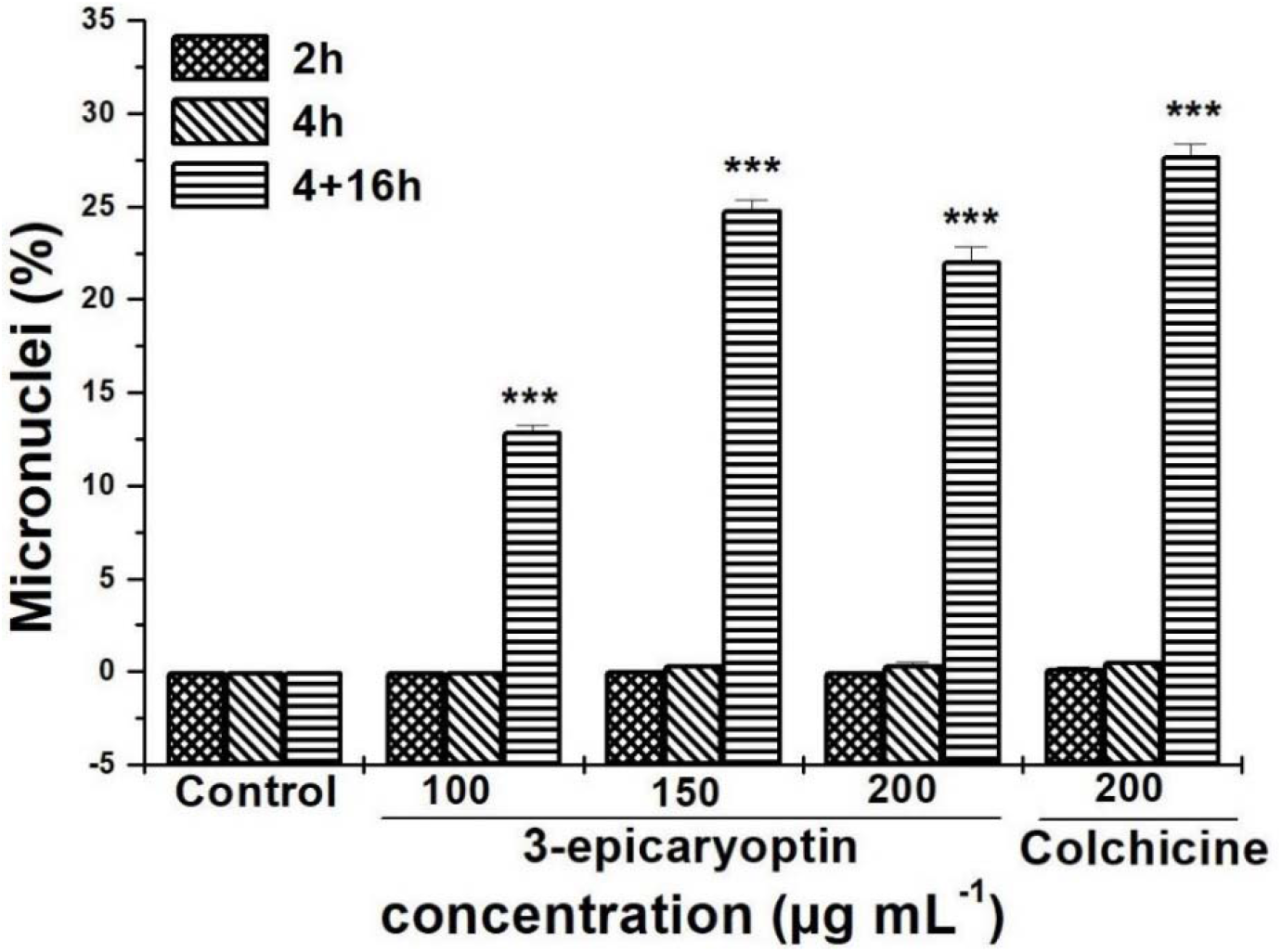
The micronuclei inducing effect of 3-epicaryoptin and colchicine on *A. cepa* root tip cells. ***Significant at *p*< 0.001 with 2 × 2 contingency χ^2^ analysis compared to respective control at *df* = 1.

**Fig. 5.**
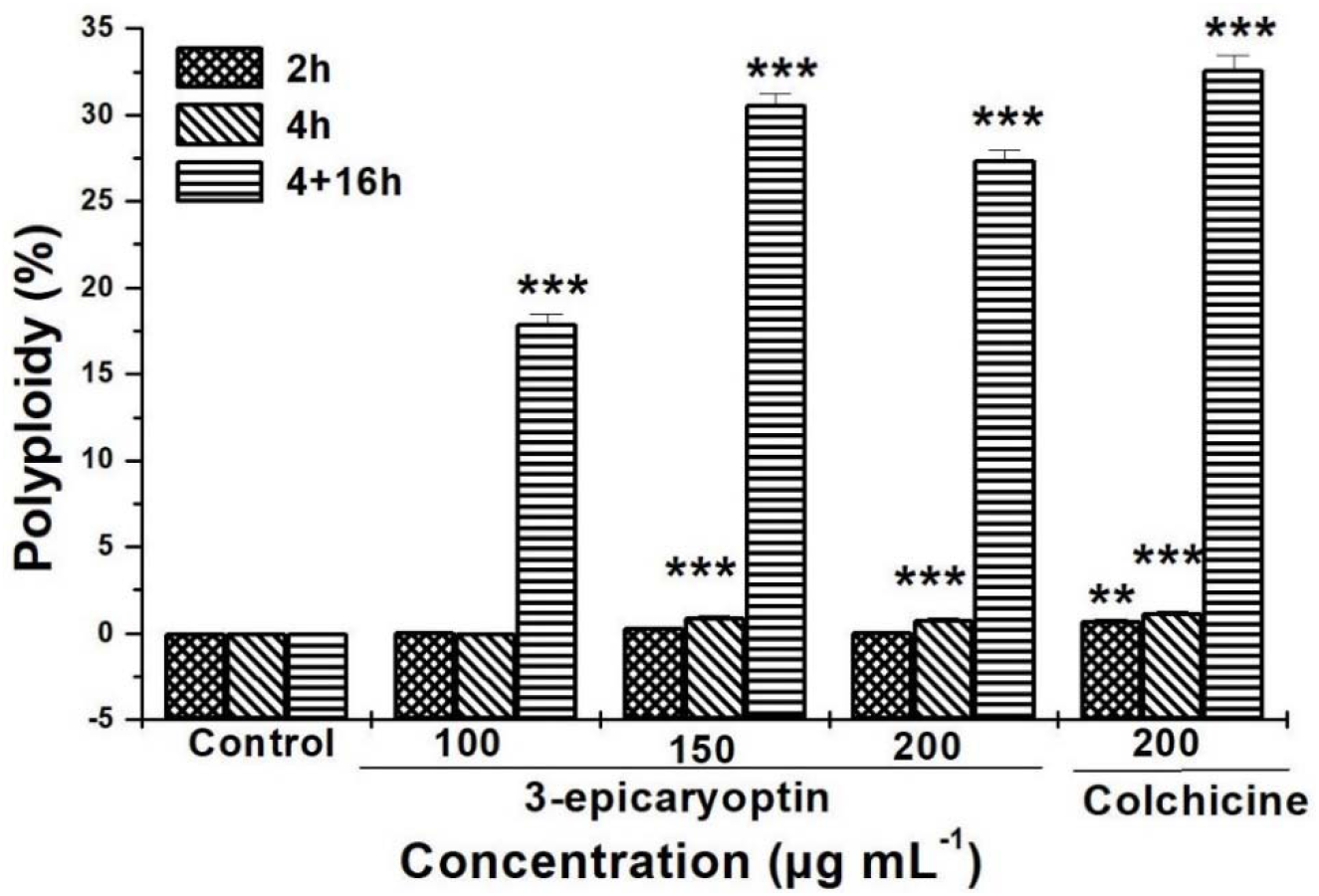
The polyploidy inducing effect of 3-epicaryoptin and colchicine on *A. cepa* root tip cells. ***Significant at *p*< 0.001, **Significant at *p*< 0.01 with 2 × 2 contingency χ^2^ analysis compared to respective control at *df* = 1.

### Correlation analysis

Pearson’s bivariate correlation analyses were performed to determine positive or negative correlation among the dividing phase and with MI. After 4 h treatment with 3-epicaryoptin, there was a strong positive correlation of metaphase with MI (*r* = 0.99227). However, prophase and anaphase had a strong negative correlation (*r* = −0.99422 and −0.99994), with MI, respectively. Prophase and anaphase also showed a strong negative correlation (*r* = −0.99986 and −0.99358) with metaphase. During 4+16 h treatment with 3-epicaryoptin, metaphase also showed a strong positive correlation (*r* = 0.91204) with MI and a strong negative correlation (*r* = −0.9484) with prophase. However, anaphase showed a low positive correlation (*r* = 0.44018) and telophase showed a strong negative correlation (*r* = −0.92074) with metaphase (**Fig. 6 & Table S5**).

**Fig. 6.**
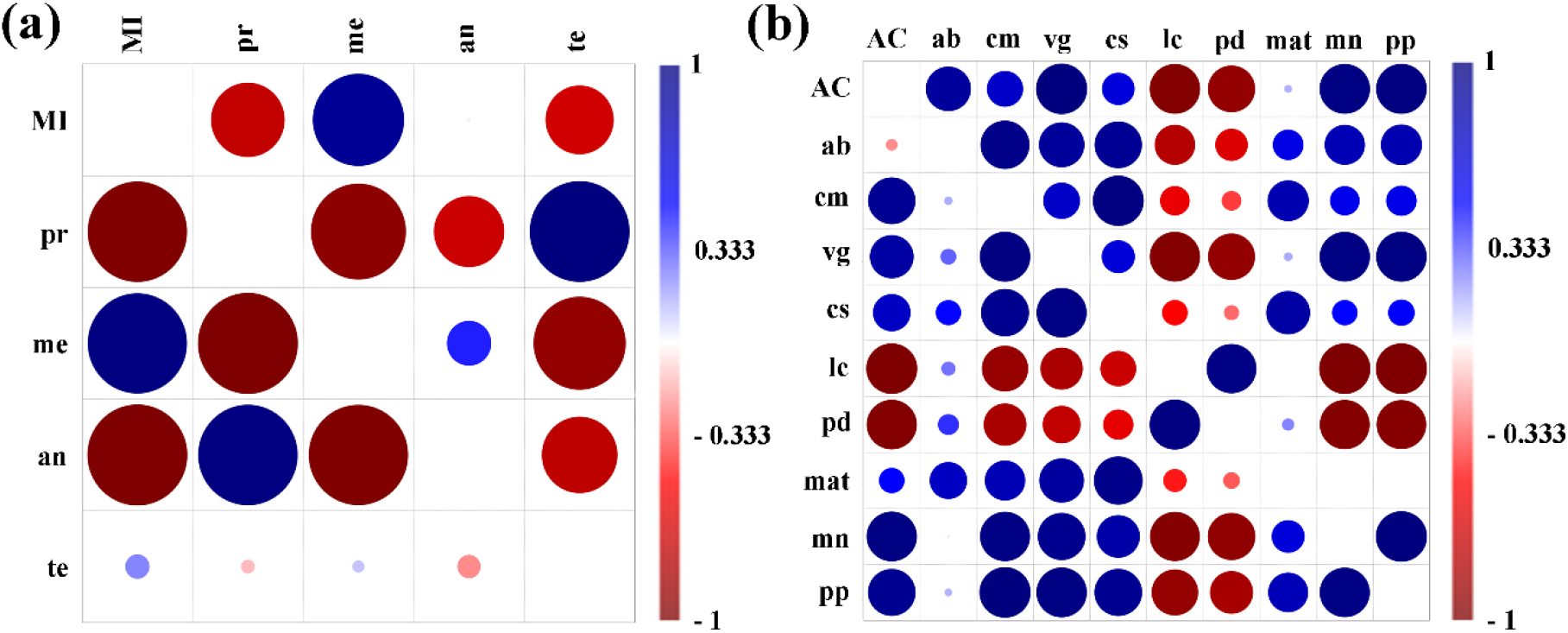
Pearson’s correlation analysis for MI, dividing phase (a), and CA (b) after 4 h and 4+16 h of water recovery treatment with 3-epicaryoptin. Here, the lower triangle showed the correlation plot for 4 h and upper triangle showed the correlation plot for 4+16 h of treatment. Mitotic index (MI), prophase (pr), metaphase (me), anaphase (an), telophase (te), aberrant cells (AC), anaphase bridge (ab), c-mitosis (cm), vagrant (vg), chromosome stickiness (cs), laggard (lc), polar deviation (pd), multipolar anaphase-telophase (mat), micronucleus (mn), and polyploid cells (pp).

Pearson’s bivariate correlation analyses was shown that after 4 h of treatment with 3-epicaryoptin, there was a strong positive correlation of c-metaphase, vagrant chromosome, MN, and PP with aberrant cells (AC) (*r* = 0.92516, 0.85639, 0.98442, and 0.93011). Similarly, vagrant, chromosome stickiness, MN, and PP were strongly correlated with c-metaphase (*r* = 0.98828, 0.93743, 0.97748, and 0.99991 respectively). There was a positive correlation of chromosome stickiness, MN, and PP with vagrant chromosome. Moreover, MN showed a strong positive correlation with PP, respectively. After 4+16 h of treatment with 3-epicaryoptin, AC showed a strong positive correlation with vagrant chromosome, PP, MN, and anaphase bridge (*r* = 0.9999, 0.98926, 0.98553, and 0.88184). While c-metaphase and chromosome stickiness had a moderately positive correlation with AC (r = 0.70739 & 0.63756). However, lagging chromosomes and polar deviation had a strong negative correlation with AC (*r* = −0.98594 and −0.92416) and was found to be similar to that of 4 h treatment. Vagrant chromosome had showed strong positive correlation (*r* = 0.98297 & 0.98704) with MN and PP cells and MN showed similar trends with PP cells (*r* = 0.99972) (**Fig. 6 and Table S6**).

## Discussion

In the present study, cytogenotoxicity of compound 3-epicaryoptin was evaluated by studying the RGR, RTS, frequencies of the different mitotic phases, MI%, and the types of structural CA as well as MN and polyploidy in *Allium cepa* test system. The *A. cepa* assay has been extremely useful for biological monitoring, investigation of environmental pollution, and determination of the toxicity of various chemical substances (Bakare et al. 2012; Bakare et al. 2013; Bakare et al. 2000; Frescura et al. 2012). The data obtained from the present study exhibited that the compound 3-epicaryoptin induced a statistically significant (*p*< 0.01 and < 0.001) and concentration-dependent *A. cepa* RGR effect with an IC_50_ value of 70.5 μg mL^-1^ at 48 h. The IC_50_ values for the RGR at 48 h are considered reliable for the assessment of the cytotoxic effects of tested compounds (Konuk et al. 2007). The RGR is the consequence of the suppression of cell division and it has been utilized for the determination of anti-proliferative effects of exogenous compounds such as pollutants, chemicals, drugs, etc. (Fiskesjo 1985). Besides the RGR effects, treatment of 3-epicaryoptin also showed RTS swelling effects (club shaped) and was identical to those of colchicine effects (**Fig. 1**). These results are consistent with previous studies, reported that the LAECI and colchicine induced *A. cepa* RTS phenomena (Barman et al. 2020; Levan 1938).

Several study reports showed that growth inhibitory effect of any chemical compound might have been manifested through its cytogenotoxic effect (Liman et al. 2018; Mercado and Caleño 2020; Leme and Marin-Morales 2009). Therefore, the present study was further extended to determine the effect of 3-epicaryoptin on the MI%, mitotic phase percentage, and CA on root meristem cells of *A. cepa*. 3-epicaryoptin (100, 150, 200 μg mL^-1^) and colchicine (200 μg mL^-1^) treatments showed an increased frequency of MI at 2 and 4 h and again decreased after 16 h of water recovery treatment as compared to control (**Table 1**). The observed increased MI% might be due to the arrest of the cell cycle in metaphase at 2 and 4 h and the later water recovery treatment for 16 h, cells revert to the interphase condition. The restitution of nuclei that they formed are PP interphase cells. This restitution causes a fall in the MI%, which was increased due to metaphase arrest during the early hours of treatment by 3-epicaryoptin and colchicine (Davidson et al. 1966). The Pearson’s bivariate correlation analysis also indicates a strong positive correlation of the increased frequency of metaphase with MI% at 4 h of treatment. We have also compared the prophase: metaphase ratio of control, 3-epicaryoptin, and colchicine treated samples. The prophase: metaphase ratio for 3-epicaryoptin (150 μg mL^-1^) was 23.22:66.20 and in the case of colchicine (200 μg mL^-1^) treated roots it was 11.3:82.35, whereas the untreated controls showed 42.58:28.84 (**Fig. 2**). This increased number of cells at metaphase in 3-epicaryoptin and colchicine treated roots may be due to the results of metaphase arrest (Davidson et al., 1966).

The results of this study further showed that 3-epicaryoptin and colchicine exposure also induced the formation of various CA like anaphase bridges, c-metaphase, vagrant chromosome, sticky chromosomes, lagging chromosome, polar deviation, and multipolar anaphase-telophase cells in the root tip cells of *A. cepa* (**Fig. 3**). Study of these chromosome abnormalities has been considered to be a promising test to determine the cytogenotoxic potential of the applied substances (Caritá and Marin-Morales 2008). The majority of the observed abnormalities induced by 3-epicaryoptin and colchicine were associated with the spindle poisoning effects such as c-metaphase, vagrant and disoriented chromosomes or chromatin dysfunction such as stickiness. Among these CAs, the highest types of abnormalities induced by 3-epicaryoptin and colchicine were c-metaphase. It may have mainly occurred due to the irregular distribution of spindle apparatus and or microtubule destabilization effects. It is the spindle poisoning effect of colchicine that shows haphazardly arranged condensed chromosomes at metaphase and therefore blocks the cell progression from metaphase to anaphase (Bonciu et al. 2018; Fiskesjö 1985; Ray et al. 2013; Barman et al. 2020). The induction of vagrant chromosomes results from the precocious movement of chromosomes in spindle poles, which leads to an unequal number of chromosome separations in the daughter nuclei and results in the formation of daughter cells with unequal sized nuclei at interphase (Bonciu et al. 2018). Chromosome stickiness is considered to be a chromatid type aberration and its occurrence indicates abnormal DNA condensation, irregular chromosome coiling, chromosome fragmentation and the formation of bridges at anaphase-telophase stages (Yüzbaşioğlu et al. 2003; Badr and Ag 1987; Klášterská et al. 1976). According to Mercykutty and Stephen (1980), stickiness may arise due to the effects of DNA depolymerization, chromatid breakage, and the stripping of the protein covering of DNA in chromosomes (Mercykutty and Stephen 1980). Besides these, 3-epicaryoptin induced other abnormalities like anaphase-bridge was formed possibly by breakage and fusion of chromatid materials, or may be due to unequal exchange of chromatid during translocation (Chatterjee and Ray 2019; Liman et al. 2010). The formation of a lagging chromosome specifies the delayed movement of the chromosome at poles and it is believed to be formed by effects of tubulin polymerization or may be due to inhibition of cytoskeletal proteins (El-Ghamery et al. 2003). The occurrence of multipolar movement of chromosome was indicative of inactivation of the spindles (Barman et al. 2021).

In addition to the CAs, cells containing MN and PP were also observed at interphase stage in all the treated concentrations (100, 150, and 200 μg mL^-1^) of 3-epicaryoptin at 4+16 h water recovery (**Fig. 3, 4, and 5**). Pearson’s correlation analysis showed a strong positive correlation between MN and PP (**Fig. 6**). The most effective concentration of 3-epicaryoptin was found to be 150 μg mL^-1^. Such type of MN cells can be originate by chromosomal break, results in the formation of acentric fragments (clastogenic action) or due to the loss of entire chromosomes as a consequence of dysfunction in normal spindle apparatus, therefore, was not incorporated with main nucleus during the course of cell division (aneugenic action) and ultimately may leads to the formation of aneuploid and PP cells in subsequent mitotic division (Chauhan et al. 1986; Chauhan and Sundararaman 1990; Fenech and Crott 2002; Yi and Meng 2003; Leme and Marin-Morales 2018). 3-epicaryoptin and colchicine induced formation of such several MN in the treated cells must probably result from its aneugenic action rather than clastogenic. On the other hand, study of PP cell frequency (30.61±0.6%, *p*< 0.001) formed by 3-epicaryoptin treatment (150 μg mL^-1^) was more or less similar to the colchicine (200 μg mL^-1^) induced PP cell percentage (32.66±0.79%, *p*< 0.001). Levan (1938) concluded that the colchicine induced formation of PP cells in *A. cepa* root tip cells occurred due to disruption in the mitotic spindle polymerization, thus preventing the migration of the chromosomes into the poles and remaining dispersed throughout the cytoplasm. At this stage, cytokinesis is not taking place and the chromatids eventually get enclosed in a new nuclear membrane and proceed into interphase as a PP cell (Levan 1938). Carvalho et al. (2019) also demonstrate that the PP cells arise when the mitotic spindle formation was hindered but the cell cycle was continued and enters into the G1 phase without completing the longitudinal chromosomes segregations, which would normally occur in anaphase and thus became PP (Carvalho et al. 2019). Thus, 3-epicaryopptin induced increased metaphase (c-metaphase) cell frequency at early hours (2 and 4 h) and the formation of MN and PP cells at 16 h recovery may be an indication that the compound 3-epicaryoptin is a spindle inhibitor similar to that of colchicine.

## Conclusions

The present study revealed that the compound 3-epicaryopptin has cytogenotoxic effects in *A. cepa*, causing root growth inhibitory and root tip swelling effects, arrest the cell division at metaphase and subsequently increase the frequency of MI. The study of cellular abnormalities specifies that the 3-epicaryoptin induced various CA, the highest occurrence was c-metaphase together with MN and PP cells. Further studies must be needed to explain the molecular mechanisms involved in the cytogenotoxicity of 3-epicaryoptin on plants.

## Acknowledgements

The authors acknowledge the DST-PURSE, DST-FIST, UGC-MRP {F.No.42-563/2013 (SR) dt. 22.3.13} and UGC-DRS-sponsored facilities in the Department of Zoology.

## Authors Contribution

**Manabendu Barman**: Investigation, statistical analysis, and draft manuscript preparation.

**Sanjib Ray**: Addressed the research problem, experiment designing, Investigation, and manuscript writing and editing.

## Availability of data and material Code availability

Mentioned in the manuscript text.

**Additional declarations for articles in life science journals that report the results of studies involving humans and/or animals. Ethics approval:** Not applicable.

