## Supplementary tables for "Cytogenotoxic effects of 3-epicaryoptin in *Allium cepa* L. root apical meristem cells"

**SUPPLEMENTARY DATA**

**Table S1.** Showing root growth retardation effects of 3-epicaryoptin in *A. cepa* root apical meristem cells. Data are means ± SE of the mean.

| Treatment | Dose  (µg mL^-1^) | Root length (cm) increase  [inhibition %] | | | IC_50_ Value  (µg mL^-1^) | | |
| --- | --- | --- | --- | --- | --- | --- | --- |
|  |  | **24h** | **48h** | **72h** | **24h** | **48h** | **72h** |
| Control | 0 | 1.88±0.09 | 3.21±0.27 | 4.03±0.15 |  |  |  |
| 3-epicaryoptin | 12.5 | 1.65±0.06  (12.11±1.36) | 2.43±0.14*  (21.75±6.9) | 3.08±0.10***  (22.96±3.98) | **68.38** | **70.5** | **63.96** |
|  | 25 | 1.46±0.09*  (21.6±5.16) | 2.2±0.05**  (28.87±6.5) | 2.76±0.1***  (30.54±4.76) |  |  |  |
|  | 50 | 1.28±0.07***  (31.15±4.94) | 2.03±0.14**  (33.22±8.91) | 2.4±0.22***  (40.92±4.7) |  |  |  |
|  | 100 | 0.55±0.04***  (70.44±2.76) | 0.63±0.07***  (79.13±3.43) | 0.63±0.06***  (83.98±2.04) |  |  |  |
|  | 200 | 0.33±0.04***  (81.89±2.65) | 0.41±0.04***  (86.82±1.24) | 0.41±0.04***  (89.77±0.85) |  |  |  |
| Colchicine | 200 | 0.16±0.04***  (90.87±2.97) | 0.21±0.04***  (92.69±2.01) | 0.28±0.05***  (92.08±0.99) |  |  |  |

*Significant at p < 0.05, **p < 0.01 and ***p < 0.001 Student’s t -test analysis compared to untreated control.

**Table S2.** Root tip swelling (at 4 h treatment + 16 h water recovery) effects of 3-epicaryoptin (100, 150 and 200 µg mL^-1^) and colchicine (200 µg mL^-1^) on *A. cepa*.

| Treatment | Concentration  (µg mL^-1^) | Roots breadth  (mm±SE) | | %  increase |
| --- | --- | --- | --- | --- |
|  |  | **Non-swollen part** | **Swollen part** |  |
| Control | 0 | 0.76±0.02 | - | - |
| 3-epicaryoptin | 100 | 0.74±0.03 | 1.44±0.05*** | 48.47 |
|  | 150 | 0.77±0.02 | 1.78±0.03*** | 56.40 |
|  | 200 | 0.71±0.03 | 1.79±0.02*** | 60.04 |
| Colchicine | 200 | 0.70±0.02 | 1.59±0.04*** | 55.92 |

*Significant at p < 0.05, **p < 0.01 and ***p < 0.001 Student’s t -test analysis compared to non-swollen part.

**Table S3.** 3-epicaryoptin and colchicine induced cell cycle delay in *A. cepa* root apical meristem cells.

| Treatment | Hours | Dose  (µg mL^-1^) | MI  (reduction %) | PM (%) | AT (%) |
| --- | --- | --- | --- | --- | --- |
| Control | 2 | 0 | 0 | 71.28±4.17 | 28.71±4.18 |
| 3-epicaryoptin |  | 100 | -45.29 | 74.07±2.31 | 25.92±2.31 |
|  |  | 150 | -116.47 | 89.27±0.59 | 10.71±0.59*** |
|  |  | 200 | -66.27 | 77.86±0.41 | 22.12±0.41 |
| Colchicine |  | 200 | -147.84 | 91.37±0.87 | 10.82±0.93*** |
| Control | 4 | 0 | 0 | 71.43±0.59 | 28.78±0.78 |
| 3-epicaryoptin |  | 100 | -37.24 | 83.03±0.59 | 16.95±0.59* |
|  |  | 150 | -109.01 | 89.43±0.87 | 10.55±0.87*** |
|  |  | 200 | -103.57 | 87.81±0.26 | 12.17±0.26*** |
| Colchicine |  | 200 | -124.14 | 93.66±1.35 | 6.28±0.73*** |
| Control | 4+16 | 0 | 0 | 65.36±0.97 | 41.82±7.89 |
| 3-epicaryoptin |  | 100 | -7.82 | 84.35±0.18 | 15.63±0.18*** |
|  |  | 150 | 4.1 | 86.67±0.86 | 13.32±0.86*** |
|  |  | 200 | 34.83 | 83.06±0.90 | 16.92±0.9** |
| Colchicine |  | 200 | 36.98 | 59.99±1.52 | 20.15±1.55** |

*Significant at *p*< 0.05, ***p*< 0.01 and ****p*< 0.001 2 × 2 contingency χ2 analysis compared to respective control with respective d*f*= 1. PM, Prophase+metaphase; AT, Anaphase+Telophase; MI, Mitotic Index.

**Table S4.** Effects of 3-epicaryoptin and colchicine on the frequency of mitotic index, micronuclei and Polyploidy induction in *A. cepa* root apical meristem cells.

| Treatment | Dose  (µg mL^-1^) | h | TC | TDC | MI (%) | Micronuclei  (%) | Polyploidy  (%) |
| --- | --- | --- | --- | --- | --- | --- | --- |
| Control | 0 | 4 h | 1287 | 71 | 5.53±0.23 | 0 | 0 |
| 3-epicaryoptin | 100 |  | 1306 | 88 | 6.76±0.57 | 0 | 0 |
|  | 150 |  | 1252 | 127 | 10.21±0.98*** | 0.36±0.02 | 0.93±0.12*** |
|  | 200 |  | 1419 | 148 | 10.42±0.41*** | 0.40±0.1 | 0.82±0.09*** |
| Colchicine | 200 |  | 1520 | 211 | 13.87±0.18*** | 0.58±0.03 | 1.22±0.04*** |
| Control | 0 | 4+16 h | 1176 | 63 | 5.34±0.17 | 0 | 0 |
| 3-epicaryoptin | 100 |  | 1161 | 64 | 5.51±0.18 | 12.94±0.34*** | 17.90±0.55*** |
|  | 150 |  | 1085 | 26 | 2.41±0.67*** | 24.84±0.54*** | 30.61±0.6*** |
|  | 200 |  | 1229 | 34 | 2.73±0.21*** | 22.10±0.73*** | 27.40±0.54*** |
| Colchicine | 200 |  | 1505 | 50 | 3.34±0.26** | 27.73±0.67*** | 32.66±0.79*** |

*Significant at *p*< 0.05, ***p*< 0.01 and ****p*< 0.001 2 × 2 contingency χ2 analysis compared to respective control with respective d*f*= 1. h, hours; TC, Total Cells; TDC; Total Dividing Cells; MI, Mitotic Index.

**Table S5.** Pearson's correlation table for mitotic abnormalities after 4 h and 4+16 h of treatment with 3-epicaryoptin.

|  | **AC** | **ab** | **cm** | **vc** | **sc** | **lc** | **pd** | **Mu** | **mi** | **po** |
| --- | --- | --- | --- | --- | --- | --- | --- | --- | --- | --- |
| **AC** |  | **0.88184** | **0.70739** | **0.9999** | **0.63756** | **-0.98594** | **-0.92416** | **0.15166** | **0.98553** | **0.98926** |
| **ab** | -0.22642 |  | **0.9571** | **0.88858** | **0.9255** | **-0.79064** | **-0.63482** | **0.59983** | **0.78915** | **0.80345** |
| **cm** | 0.92516 | 0.16024 |  | **0.71755** | **0.99554** | **-0.57932** | **-0.38372** | **0.80593** | **0.57734** | **0.59648** |
| **vc** | 0.85639 | 0.30902 | 0.98828 |  | **0.64864** | **-0.98342** | **-0.91853** | **0.16596** | **0.98297** | **0.98704** |
| **sc** | 0.73511 | 0.49389 | 0.93743 | 0.97959 |  | **-0.49985** | **-0.2949** | **0.85819** | **0.49774** | **0.51811** |
| **lc** | -0.99872 | 0.27549 | -0.90474 | -0.82912 | -0.69982 |  | **0.975** | **0.01565** | **-1** | **-0.99977** |
| **pd** | -0.98148 | 0.40883 | -0.83531 | -0.74161 | -0.59162 | 0.98992 |  | **0.23744** | **-0.97554** | **-0.97007** |
| **Mu** | 0.49912 | 0.73102 | 0.79068 | 0.87486 | 0.95437 | -0.45457 | -0.32386 |  | **-0.01808** | **0.005566** |
| **mi** | 0.98442 | -0.05165 | 0.97748 | 0.93383 | 0.84286 | -0.97425 | -0.93251 | 0.64369 |  | **0.99972** |
| **po** | 0.93011 | 0.14715 | 0.99991 | 0.98617 | 0.93273 | -0.9103 | -0.84252 | 0.7825 | 0.98019 |  |

Here, the black color showed the correlation matrix for 4 h and red color showed the correlation matrix for 4+16 h of treatment. Aberrant cells (AC), anaphase bridge (ab), c-mitosis (cm), vagrant (vg), chromosome stickiness (cs), laggard (lc), polar deviation (pd), multipolar anaphase-telophase (mat), micronucleus (mi), and polyploid cells (po).

**Table S6.** Pearson's correlation table for MI and dividing phase after 4 h and 4+16 h of treatment with 3-epicaryoptin.

|  | **MI** | **pr** | **me** | **an** | **te** |
| --- | --- | --- | --- | --- | --- |
| **MI** |  | **-0.73494** | **0.91204** | **0.033231** | **-0.67975** |
| **pr** | -0.99422 |  | **-0.9484** | **-0.70218** | **0.99695** |
| **me** | 0.99227 | -0.99986 |  | **0.44018** | **-0.92074** |
| **an** | -0.99994 | 0.99535 | -0.99358 |  | **-0.75563** |
| **te** | 0.23776 | -0.13211 | 0.11537 | -0.22698 |  |

Here, the black color showed the correlation matrix for 4 h and red color showed the correlation matrix for 4+16 h of treatment. Mitotic index (MI), prophase (pr), metaphase (me), anaphase (an), telophase (te),
